# The contribution of astrocyte and neuronal Panx1 to seizures is model and brain region dependent

**DOI:** 10.1101/2021.01.12.426355

**Authors:** Price Obot, Libor Velíšek, Jana Velíšková, Eliana Scemes

## Abstract

Pannexin1 (Panx1) is an ATP release channel expressed in neurons and astrocytes that plays important roles in CNS physiology and pathology. Evidence for the involvement of Panx1 in seizures includes the reduction of epileptiform activity and ictal discharges following Panx1 channel blockade or deletion. However, very little is known about the relative contribution of astrocyte and neuronal Panx1 channels to hyperexcitability. To this end, mice with global and cell type specific deletion of Panx1 were used in one *in vivo* and two *in vitro* seizure models. In the low-Mg^2+^ *in vitro* model, global deletion but not cell-type specific deletion of Panx1 reduced the frequency of epileptiform discharges. This reduced frequency of discharges did not impact the overall power spectra obtained from local field potentials. In the *in vitro* KA model, in contrast, global or cell type specific deletion of Panx1 did not affect the frequency of discharges, but reduced the overall power spectra. EEG recordings following KA-injection *in vivo* revealed that although global deletion of Panx1 did not affect the onset of status epilepticus (SE), SE onset was delayed in mice lacking neuronal Panx1 and accelerated in mice lacking astrocyte Panx1. EEG power spectral analysis disclosed a Panx1-dependent cortical region effect; while in the occipital region, overall spectral power was reduced in all three Panx1 genotypes; in the frontal cortex, the overall power was not affected by deletion of Panx1. Together, our results show that the contribution of Panx1 to ictal activity is model, cell-type and brain region dependent.

## Introduction

Epilepsy is a burden worldwide and, in the U.S., represents the 4^th^ most common chronic neurologic disorder, afflicting more than 1% of the population ((IOM). 2012; England et al., 2012; Zack and Kobau, 2017). A widely accepted view in the field is that epileptic seizures largely relate to an imbalance in brain excitatory and inhibitory tone mainly contributed by glutamate and GABA release from neurons. However, the contribution of other transmitters and modulators such the purinergic and adenosinergic signaling is gaining substantial support in epilepsy research. Early evidence for the excitatory effect of purinergic signaling includes augmentation of seizures in limbic structures by intracerebroventricular ATP injection in a mouse model with focal onset of status epilepticus (Engel et al., 2012; Engel et al., 2016) and that ATP sensitive P2X receptors promote depolarizing currents in CA1 and CA3 hippocampal pyramidal neurons. On the other hand, the inhibitory effect of adenosine, a byproduct of ATP hydrolysis, has been long known to have anti-convulsant effects through its action on A1 receptors (Barraco et al., 1984; O’Shaughnessy et al., 1988).

Both neurons and astrocytes can release ATP via regulated secretion (Coco et al., 2003; Pankratov et al., 2006; Bowser and Khakh, 2007) or diffusion through ion channels, such as CALHM and hemichannels (connexins and pannexins) (Siebert et al., 2013; Retamal et al., 2014). Among these channels, a substantial number of studies indicate that pannexin1 (Panx1) contributes to epileptic seizures (Thompson et al., 2008; Santiago et al., 2011; Dossi et al., 2018). For instance, Panx1 has been shown to promote epileptiform-like activity in hippocampal slices following N-methyl-D-aspartic acid (NMDA) receptor activation by providing the secondary NMDA depolarizing current that increases burst activity (Thompson et al., 2008). Using the kainic acid (KA)-seizure model in juvenile mice, our laboratory provided the first *in vivo* evidence that Panx1 channels contribute to the maintenance of status epilepticus (SE) by releasing ATP (Santiago et al., 2011). In addition, evidence was recently provided that Panx1 plays a role in seizure initiation by releasing ATP in epileptic human cortical brain tissue (Dossi et al., 2018).

The recognition for an astrocytic basis of epilepsy includes the modulation of synaptic strength and plasticity by astrocyte-released ATP acting on post-synaptic P2X receptors (Boué-Grabot and Pankratov, 2017), and the regulation of basal extracellular adenosine levels and seizure activity by an astrocytic enzyme, adenosine kinase (ADK) (Boison, 2010). In this regard, our recent study evaluating the relative contribution of astrocyte and neuronal Panx1 to KA seizures indicated that the faster SE onset recorded from mice lacking astrocyte Panx1 compared to that of mice lacking neuronal Panx1 was due to increased astrocyte ADK levels in the former (Scemes et al., 2019). To further evaluate the impact of neuronal and astrocyte Panx1 to brain electrical activity during seizures, we used two *in vitro* seizure models (triggered by exposure to low-Mg^2+^ or KA conditions) and performed EEG recordings in KA-injected mice lacking Panx1 in neurons or astrocytes. Our results indicate that both neuronal and astrocyte Panx1 contribute to seizures and that their relative contribution is different depending on both model and brain region that is studied.

## Materials and Methods

### Ethics statement

All procedures described below were reviewed and approved by the Institutional Animal Care and Use Committee (IACUC) of New York Medical College (NYMC). The procedures follow ARRIVE guidelines and are consistent with the Guide for the Care and Use of Laboratory Animals, 8^th^ Edition. Mice colonies were housed and maintained in NYMC’s AALAC-accredited Laboratory Animal Resource Facility with a 12 hr light/dark cycle (lights on at 7:00 am) and with access to food and water *ad-libitum*. Mice used were generated as previously described (Hanstein et al., 2013; Scemes et al., 2019), and all experiments were performed on three weeks (P21-25) old male and female Panx1^f/f^ (control), Panx1^tm1a(KOMP)Wtsi^ (global Panx1-null), mGFAP-Cre:Panx1^f/f^ (astrocyte Panx1-null), and mNFH-Cre:Panx1^f/f^ (neuronal Panx1-null) mice raised on C57BL/6 background.

### Slice preparation

Hippocampal slices were prepared as previously described (Kirchner et al., 2006; Velísek et al., 2000). Briefly, brains were rapidly removed and immediately submerged in ice-cold (3-4 °C) artificial cerebrospinal fluid (aCSF) containing (in mM): 126 NaCl, 5 KCl, 26 NaHCO_3_, 2 CaCl_2_, 2 MgSO_4_, 1.25 NaH_2_PO_3_, and 10 D-glucose; pH 7.4 ± 0.1. Hippocampal slices (400 μm thick) containing the entorhinal-hippocampal connections were cut approximating the horizontal plane using a vibratome (Leica VT1000 S) and transferred to an interface-type chamber and perfused with aCSF (~2.5 ml/min) maintained at 34 ± 1 °C, bubbled with 95% O_2_ and 5% CO_2_ for at least one hour recovery before recordings started.

### Field potential recordings

The viability of slices was first tested by examining potentials evoked by stimulation of Schaffer collaterals using a bipolar stimulating electrode and recording in the stratum pyramidale of CA1 regions of the hippocampi by extracellular glass micropipette electrodes filled with 2 M NaCl. After 5 min of field potential recording under normal aCSF (baseline), solution was switched to Mg^2+^-free aCSF for >80 min or to regular aCSF containing 1 μM kainic acid (KA) for 30 min. AxoScope 10.3 software was used in data acquisition and Clampfit 11.0.3 (Molecular Devices) was used to evaluate the onset, duration, number and frequency of induced epileptiform activity.

### Electroencephalographic (EEG) recordings

EEG activity was recorded as previously described in (Scemes et al., 2019). Briefly, electrodes were implanted in 3-week old mice using the stereotactic apparatus (Heinrich Kopf Inc.). Deep anesthesia was induced with isoflurane (5% of isoflurane in O_2_ for induction in an induction chamber and 2% isoflurane in O_2_ for maintenance through the mask attached to the stereotactic apparatus). The depth of anesthesia was verified by absence of corneal and toe-pinch reflexes assessed every 3 mins. Skin overlying the skull was retracted to expose skull surface, and stainless-steel screws (PlasticsOne) were placed behind lambda and in the nasal bone for ground and reference electrodes, respectively. Silver ball electrodes for cortical recordings were placed epidurally over the frontal and occipital cortices, bilaterally and symmetrically, and connected to an acquisition computer via dual-in-line connectors and preamplifiers. The electrodes were secured, and a cap was built with dental acrylic. After surgery, mice were placed on a heating pad until full ambulation. Two days after surgery, mice were injected with 20 mg/kg of kainic acid (KA) i.p. and EEG recordings monitored through Sirenia EEG/video monitoring system (Pinnacle Technology). Ictal activity was determined by EEG recordings in combination with continuous video monitoring of behavioral seizures. EEG ictal activity was defined as periods of rhythmic high amplitude spike and wave discharges with a minimum duration of 5 sec. accompanied initially by a sudden arrest of spontaneous behavior (i.e., grooming, walking, exploratory ambulation, etc.), head bobbing, myoclonic twitches progressing into forelimb clonus (unilateral or bilateral with/without rearing). The end of a seizure was noted as recovery of baseline EEG activity and re-gaining of spontaneous behavior. This differs from SE, which was determined to begin with seizure that did not show behavioral recovery even during brief periods of suppressed EEG activity. For the progression of seizures, the time interval between the onset of first spike to the first EEG seizure and to the onset of SE were evaluated.

### Power spectra analysis

Power spectra of EEG ictal activity was analyzed by the Score Frequency Analysis (SFA) function in Sirenia Sleep Pro 1.7.4. For power spectra analysis of local field potentials, epileptiform activity was first converted into EEG signals using File Format Converter (National Instruments) before SFA. To avoid the contribution of noise to the power spectra analysis, only local field potentials with high signal/noise ratio were used. Using Fast Fourier Transform, EEG and local field potential frequency distributions were assessed in 10 s epochs after band-pass filtering at 1 Hz (high pass) and 100 Hz (low pass). The overall changes in power were performed using the area under the curves obtained from the averaged frequency-power distribution using GraphPad Prism 9.0. When appropriate, the frequency-power distribution for delta (0.5 – 5.0 Hz), theta (>5 – 8 Hz), alpha (>8 – 13 Hz), beta (>13 – 30 Hz), and gamma (>30 – 50 Hz) bands was also determined.

### Statistical analyses

Data are expressed as mean ± s.e.m. Depending on the experimental design, unpaired t-test, one-way or two-way ANOVA with repeated measures followed by Dunnett’ or Tukey’ multiple comparison tests were performed using GraphPad Prism 9.0, as specified in Results. Statistical significance was considered at p < 0.05.

## Results

### 1) Deletion of Panx1 from astrocytes or neurons is not sufficient to reduce low Mg^2+^-induced epileptiform discharges in vitro

To evaluate the contribution of astrocyte and neuronal Panx1 channels to hippocampal hyperactivity, we used Mg^2+^-free aCSF, an established model of epileptiform activity (Velíšková and Velíšek, 2013; Kirchner et al., 2006; Engel et al., 2000; Velísek et al., 1994) that has been previously used to disclose the pro-ictal effects of Panx1 channels (Thompson et al., 2008; Dossi et al., 2018). To first validate previous reports showing that Panx1 contributes to epileptiform discharges, we used hippocampal slices of Panx1^f/f^ and Panx1-null mice. Following Mg^2+^-free aCSF application, initial epileptiform activity of positive and negative polarities occurred within 13-16 min in both genotypes (Fig. 1A) and continued upon bathing in Mg^2+^-free aCSF, subsiding quickly following switch to normal aCSF (Supplemental Figure S1). In the presence of low-Mg^2+^ aCSF, the number of epileptiform discharges in both genotypes increased over time reaching a plateau around 45 – 50 min (Fig. 1B). About 70% of slices irrespective of a genotype exhibited large, negative shifts in field potentials (−10 to −20 mV) characteristic of spreading depolarizations (SDs) (Supplemental Figure S2). No significant differences in terms of the latency to first discharge from onset of perfusion with low-Mg^2+^ aCSF or the duration of epileptiform activity were recorded between Panx1^f/f^ and Panx1-null slices (Table 1). The ictal-like activity in slices from Panx1^f/f^ appeared as burst-like discharges interrupted by periods of baseline activity (≥ 60 sec) (Fig. 1A, top trace). In contrast, Panx1-null slices did not show the burst-like activity but rather exhibited continuous epileptiform discharges (Fig. 1A, bottom trace). The frequency of discharges recorded from Panx1-null (16.52 ± 4.39 discharges/min) was significantly lower than that of Panx1^f/f^ slices (58.02 ± 10.40 discharges/min; ANOVA followed by Dunnett’s test, p = 0.009; Fig. 1D). Thus, these data confirm previous reports indicating that Panx1 enhances the frequency of epileptiform discharges.

**FIGURE 1.**
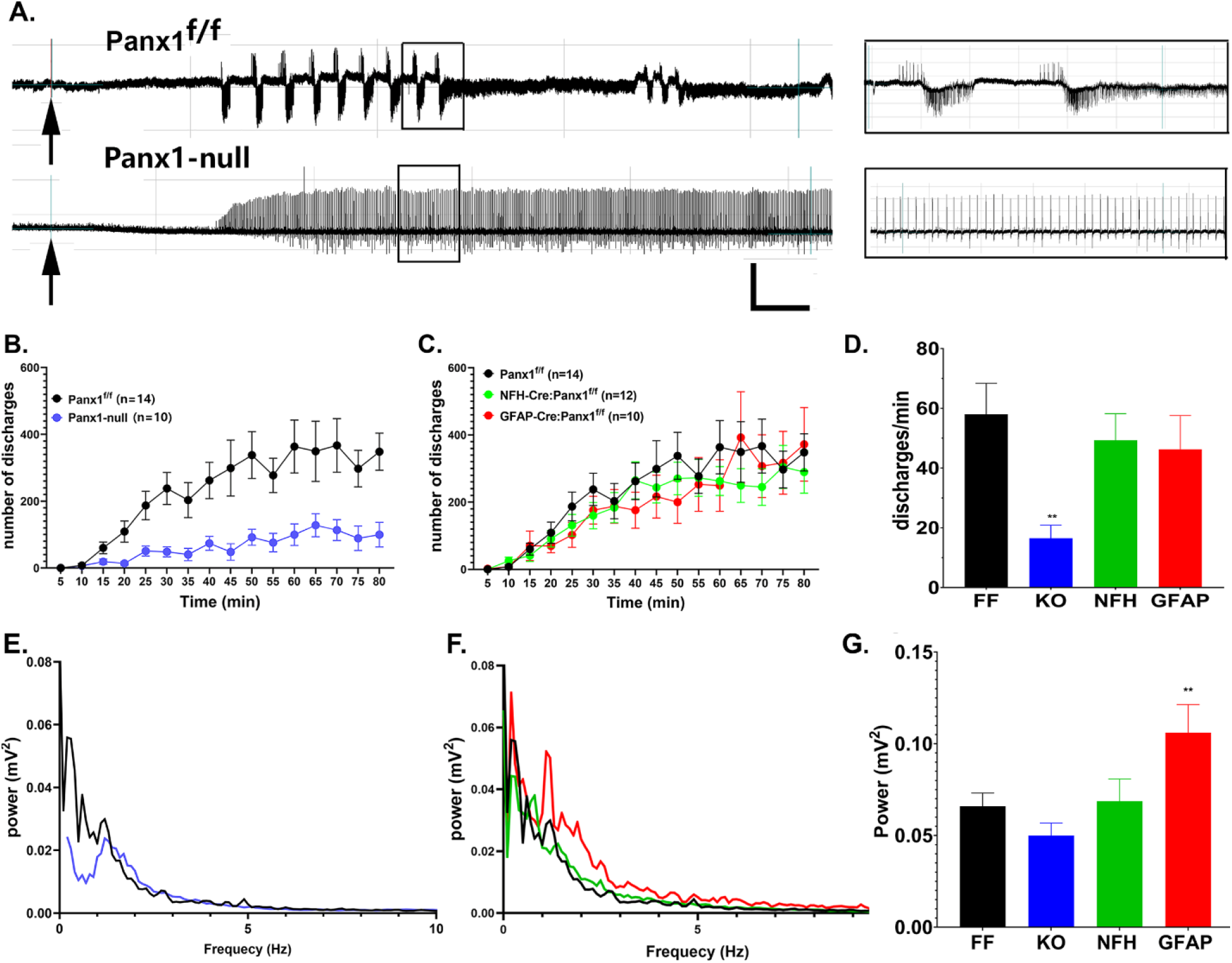
Effects of Panx1 deletion on epileptiform activity induced by low Mg^2+^condition. (**A**) Representative extracellular field potential recordings obtained from CA1 regions of Panx1^f/f^ (top) and Panx1-null (bottom) hippocampal slices upon exposure to low-Mg^2+^ aCSF (arrows). Calibration bars: 5.0 mV (vertical), 5 min (horizontal). *Insets*: Five min expansions of boxed recording segments. Note the burst-like epileptiform activity pattern of Panx1^f/f^ slices and the non-burst discharge pattern in Panx1-null slices. (**B-C**) Mean ± SEM values of the number of discharges (computed in 5 min bins) over time recorded upon Mg^2+^-free aCSF exposure of (**B**) Panx1^f/f^ (black symbols) and Panx1-null (blue symbols) slices, and (**C**) from NFHCre:Panx1^f/f^ (green symbols) and GFAPCre:Panx1^f/f^ (red symbols) hippocampal slices. (**D**) Mean ± s.e.m. values of frequency of discharges (discharges/min) recorded from Panx1^f/f^ (FF), Panx1-null (KO), NFH-Cre:Panx1^f/f^ (NFH), and GFAP-Cre:Panx1^f/f^ (GFAP) hippocampal slices; frequency values were obtained by dividing the total number of discharges by total duration of epileptiform activity. (**E-F**) Power spectra obtained from hippocampal slices of (**E**) Panx1^f/f^ and Panx1-null, and (**F**) Panx1^f/f^, NFH-Cre:Panx1^f/f^ and GFAP-Cre:Panx1^f/f^ mice. (**G**) Mean ± s.e.m. values of the overall power recorded from Panx1^f/f^ (FF), Panx1-null (KO), NFH-Cre:Panx1^f/f^, and GFAP-Cre:Panx1^f/f^ (GFAP) hippocampal slices obtained from the areas under the curves displayed in parts **E** and **F**. The averaged power spectra and the overall power measures were determined within 0 – 10 Hz range. N = number of slices. P values calculated from one-way ANOVA followed by Dunnett’s multiple comparison tests (*p<0.05; **p<0.001).

**Table 1.** Epileptiform activity parameters recorded under low-Mg^2+^ aCSF. Mean ± s.e.m. values of the time to first spike onset (onset), duration of epileptiform activity (DEA), and total number of discharges obtained in hippocampal slices of Panx1^f/f^ (FF), Panx1-null (KO), NFH-Cre:Panx1^f/f^ (NFH), and GFAP-Cre:Panx1^f/f^ (GFAP) mice. Onset: time interval between bath-application of Mg^2+^-free aCSF and the first spike; DEA: time interval between the first spike and washout of Mg^2+^-free aCSF; Discharges: total number of discharges recorded during DEA. N = number of hippocampal slices. P values were obtained from one-way ANOVA, followed by Tukey’s multiple comparisons (**p<0.001, compared to FF).

|  | **FF**<br>(n=14) | **KO**<br>(n=10) | **NFH**<br>(n=12) | **GFAP**<br>(n=10) | **ANOVA** |
| --- | --- | --- | --- | --- | --- |
| **Onset**<br>(min) | 13.6±1.0 | 13.9±2.0 | 15.8±2.4 | 14.1±1.3 | F(3,42)=0.34<br>$p=0.80$ |
| **DEA**<br>(min) | 66.4±1.0 | 65.1±2.3 | 63.8±2.3 | 65.9±2.3 | F(3,42)=0.44<br>$p=0.73$ |
| **Discharges**<br>(number) | 3813.0±662.6 | 1051.0±259.6** | 3036.0±494.2 | 3101.0±810.7 | F(3,42)=3.66<br>**$p=0.02$** |

To evaluate the relative contribution of neuronal and astrocyte Panx1 to the epileptiform activity, we used slices from mice lacking Panx1 in neurons or in astrocytes. The pattern of epileptiform discharges under low-Mg^2+^ condition in slices from NFH-Cre:Panx1^f/f^ and GFAP-Cre:Panx1^f/f^ mice was similar to that recorded from Panx1^f/f^ (Supplemental Figure S3), and no significant differences in terms of latency to first discharge and duration of epileptiform activity were recorded between the three genotypes (Table 1). As per the time course of epileptiform discharges, neither the progression (Fig. 1C) nor the frequency of discharges (Fig. 1D) differed among the three genotypes (ANOVA followed by Dunnett’ test, p > 0.05). This surprising result indicates that neither deletion of Panx1 from astrocytes nor from neurons are individually sufficient to decrease epileptiform discharges seen in the global Panx1-null slices.

Despite the reduced frequency of discharges recorded from Panx1-null slices (Fig 1B) and lower number of discharges (Table 1), statistical analysis of the overall power spectra indicated no significant differences between Panx1-null slices (0.050 ± 0.010 mV^2^) and Panx1^f/f^ (0.066 ± 0.007 mV^2^; ANOVA followed by Dunnett’s test, p = 0.528) (Fig. 1G). This apparent discrepancy was due to increased amplitudes of epileptiform discharges in Panx1-null slices (4.34 ± 0.28 mV) compared to those recorded from Panx1^f/f^ slices (2.75 ± 0.45 mV; t-test, p=0.012) (not shown). In contrast to Panx1-null, a significant difference in power spectra was detected between Panx1^f/f^ slices and those from mice lacking astrocyte Panx1 (Fig. 1G); the overall power spectra in GFAP-Cre:Panx1^f/f^ slices (0.106 ± 0.015 mV^2^) was significantly higher than that of Panx1^f/f^ slices (0.066 ± 0.007 mV^2^; ANOVA followed by Dunnett’s test, p = 0.027), which was due to increased delta (0.5-5 Hz) band activity (Fig. 1F). Power spectra of NFH-Cre:Panx1^f/f^ (0.069 ± 0.012 mV^2^) did not differ from that of Panx1^f/f^ slices (ANOVA followed by Dunnett’s test, p = 0.999).

Together, these results indicate that although global deletion of Panx1 leads to a reduction in the frequency of epileptiform discharges, this reduction was not sufficient to significantly alter the overall power spectra of epileptiform activity recorded from hippocampi of these mice. In addition, because neither deletion of neuronal nor astrocyte Panx1 was sufficient to reduce the frequency of epileptiform discharges, it is likely that Panx1 channels in both cell type populations are necessary to sustain the epileptiform discharges under the low-Mg^2+^ model.

### 2) Astrocyte and neuronal Panx1 participate to increase KA-induced epileptiform discharges

Given the frequent use of KA to induce seizures and epileptiform activity (Lothman et al., 1981; Fisher and Alger, 1984; Ben-Ari and Cossart, 2000) and the high levels of AMPA/kainate receptors in the rodent hippocampus (Monaghan and Cotman, 1982; Wisden and Seeburg, 1993; Paternain et al., 2000), we used this model to further evaluate the contribution of Panx1 to epileptiform activity. Bath application of 1 μM KA to hippocampal slices from Panx1 transgenic mice generated epileptiform discharges that were positive and/or negative in polarity; all genotypes displayed initial burst-like discharges that were often followed by continuous non-burst-like discharges of various durations (but no more than 25 min), which were genotype dependent (Figs. 2 A-D). Interestingly, this epileptiform activity resumed upon washout of KA with normal aCSF (Supplemental Figure S4). No significant differences in terms of the latency to first discharge and the duration of epileptiform activity were recorded between Panx1^f/f^ and Panx1-null slices exposed to KA (Table 2). As shown in Figure 2E, the number of epileptiform discharges recorded from Panx1^f/f^ and Panx1-null slices quantified within 5 min bins increased over time reaching a peak around 15 min and subsiding 25-30 min after KA application. The frequency of discharges recorded from Panx1-null slices although reduced (48.05 ± 7.75 discharges/min) was not significantly different from that of Panx1^f/f^ slices (71.97 ± 10.89 discharges/min; ANOVA followed by Tukey’s test, p = 0.96; Fig. 2G). Thus, differently from what we found for the low-Mg^2+^ model (Fig. 1), global deletion of Panx1 does not significantly attenuate the frequency of epileptiform discharges induced by KA.

**FIGURE 2.**
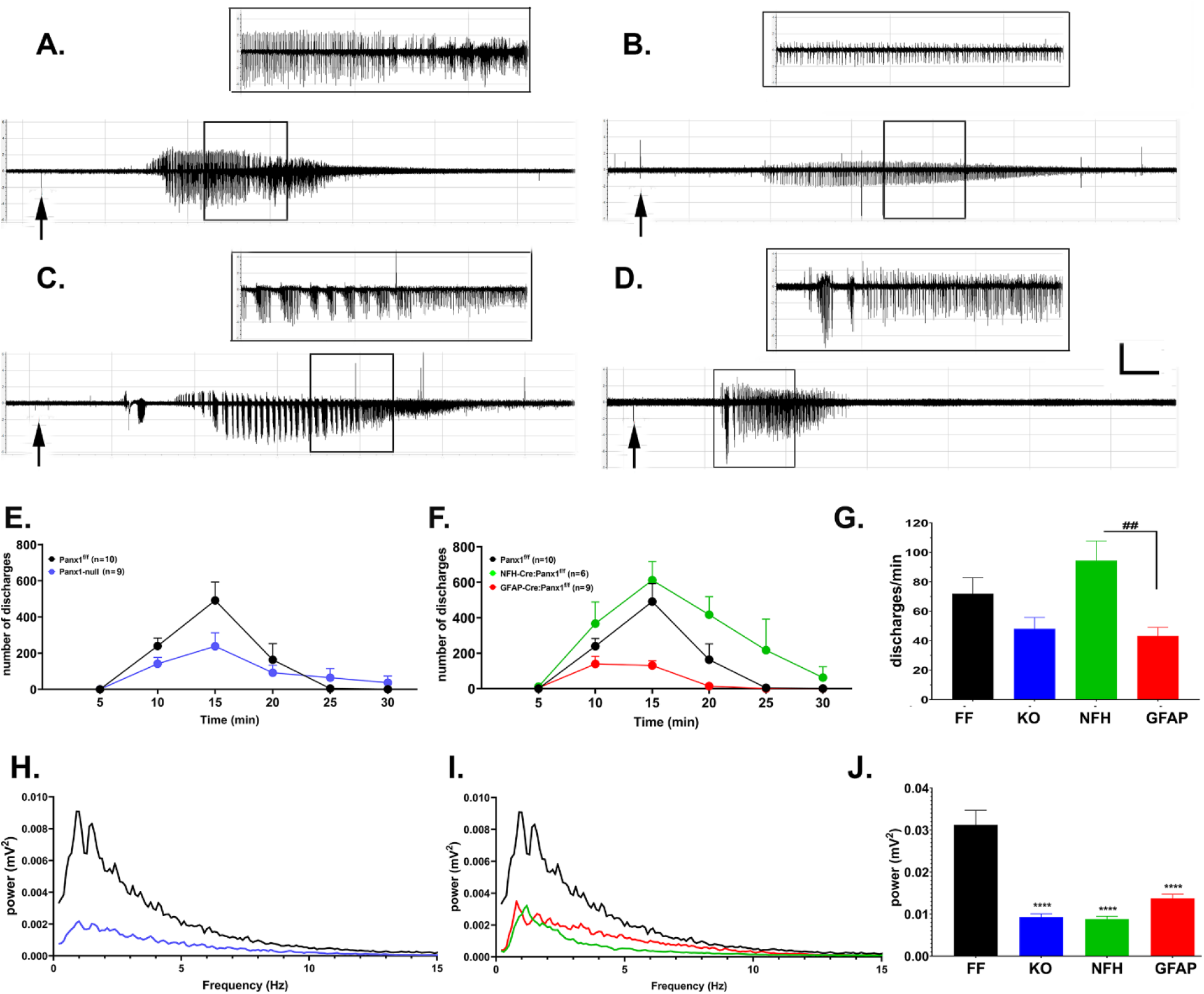
Contribution of astrocyte and neuronal Panx1 to KA-induced epileptiform discharges. (**A-D**) Representative extracellular field potential recordings obtained from CA1 region of Panx1^f/f^ (**A**), Panx1-null (**B**), NFH-Cre:Panx1^f/f^ (**C**), and GFAP-Cre:Panx1^f/f^ (**D**) mice hippocampal slices following 1 μM KA exposure (arrows). Calibration bars: 2.0 mV (vertical), 2.5 min (horizontal). *Insets*: 5-min expansions of boxed recording segments. (**E-F**) Time courses of discharges recorded from (**E**) Panx1^f/f^ (black symbols) and Panx1-null (blue symbols), and (**F**) from NFH-Cre:Panx1^f/f^ (green symbols) and GFAP-Cre:Panx1^f/f^ (red symbols) hippocampal slices. The mean ± s.e.m. values of the number of discharges recorded from the slices were computed in 5 min bins starting upon KA perfusion. (**G**) Mean ± s.e.m. values of frequency of discharges (discharges/min) recorded from Panx1^f/f^, NFH-Cre:Panx1^f/f^, GFAP-Cre:Panx1^f/f^, and Panx1-null hippocampal slices obtained by dividing the total number of discharges by the duration of epileptiform activity. One-way ANOVA followed by Tukey’s multiple comparison test. ##p<0.001. (**H-I**) Power spectra obtained from hippocampal slices of Panx1^f/f^ and Panx1-null (**H**), and from and Panx1^f/f^, NFH-Cre:Panx1^f/f^ and GFAP-Cre:Panx1^f/f^ mice (**I**). (**J**) Mean ± s.e.m. values of the overall power recorded from Panx1^f/f^ (FF), Panx1-null (KO), NFH-Cre:Panx1^f/f^ (NFH), and GFAP-Cre:Panx1^f/f^ (GFAP) hippocampal slices; overall power was obtained from the areas under the curves displayed in parts **H** and **I**. The average power spectra and the overall power measures were determined within 0 – 15 Hz frequency range. N = number of slices. One-way ANOVA followed by Dunnett’s multiple comparison test. ****P<0.0001.

**Table 2.**
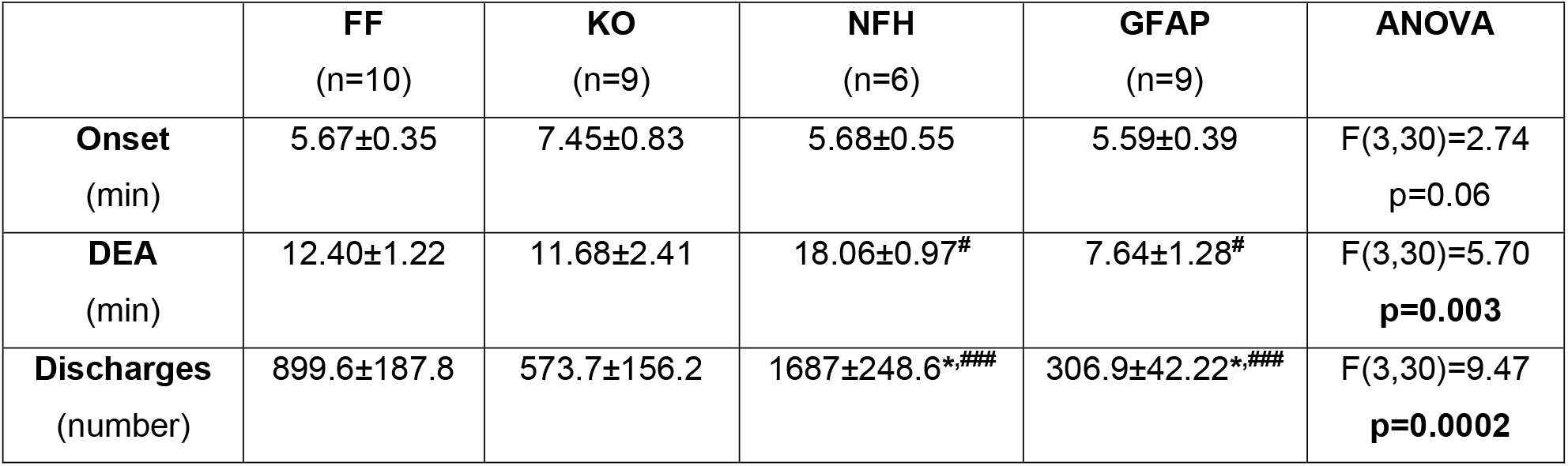
Epileptiform activity parameters recorded under KA-condition. Mean ± s.e.m. values of the onset time to first spike (onset), duration of epileptiform activity (DEA), and total number of discharges obtained in hippocampal slices of Panx1^f/f^ (FF), Panx1-null (KO), NFH-Cre:Panx1^f/f^ (NFH), and GFAP-Cre:Panx1^f/f^ (GFAP) mice. Onset: time interval between bath-application of 1 μM KA and the first spike; DEA: time interval between the first spike and the end of epileptiform activity; Discharges: total number of discharges recorded during DEA. N = number of hippocampal slices. P values were obtained from one-way ANOVA, followed by Tukey’s multiple comparisons (* p<0.05 compared to FF; ^#^p<0.05, ^###^p<0.001 for NFH x GFAP).

|  | **FF**<br>(n=10) | **KO**<br>(n=9) | **NfH**<br>(n=6) | **GFAP**<br>(n=9) | **ANOVA** |
| --- | --- | --- | --- | --- | --- |
| **Onset**<br>(min) | 5.67±0.35 | 7.45±0.83 | 5.68±0.55 | 5.59±0.39 | F(3,30)=2.74<br>$p=0.06$ |
| **DEA**<br>(min) | 12.40±1.22 | 11.68±2.41 | 18.06±0.97<sup>#</sup> | 7.64±1.28<sup>#</sup> | F(3,30)=5.70<br>**$p=0.003$** |
| **Discharges**<br>(number) | 899.6±187.8 | 573.7±156.2 | 1687±248.6<sup>*,###</sup> | 306.9±42.22<sup>*,###</sup> | F(3,30)=9.47<br>**$p=0.0002$** |

Regarding the contribution of astrocyte and neuronal Panx1 to KA-induced epileptiform discharges, no significant differences in terms of onset of first discharge was detected among the Panx1^f/f^, GFAP-Cre:Panx1^f/f^ and NFH-Cre:Panx1^f/f^ (Table 2). In terms of duration of epileptiform activity, a significant difference was recorded only between NFH-Cre:Panx1^f/f^ and GFAP-Cre:Panx1^f/f^ slices, with NFH-Cre:Panx1^f/f^ slices showing longer (18.06 ± 0.97 min) and GFAP-Cre:Panx1^f/f^ shorter durations (7.64 ± 1.28 min; Table 2). A significant difference in the total number of discharges was recorded between Panx1^f/f^, NFH-Cre:Panx1^f/f^, and GFAP-Cre:Panx1^f/f^ slices; in this case NFH-Cre:Panx1^f/f^ slices showed increased (1687.0 ± 248.6 discharges) while GFAP-Cre:Panx1^f/f^ slices showed reduced number of discharges (306.9 ± 42.2 discharges) compared to Panx1^f/f^ (899.6 ± 187.8 discharges; p = 0.0002; Table 2). The frequency of discharges recorded from slices lacking astrocyte Panx1 (43.11 ± 5.92 discharges/min) was significantly lower than that recorded from slices lacking neuronal Panx1 (94.41 ± 13.34 discharges/min; ANOVA followed by Tukey’s test, p = 0.002); however, neither significantly differed from that recorded from Panx1^f/f^ (71.97 ± 10.89 discharges/min; Fig. 2G).

These data thus suggest that differently from the low-Mg^2+^ model, global deletion of Panx1 does not affect the frequency of discharges following exposure to KA. Such lack of effect of global deletion of Panx1 on the frequency of discharges is likely resultant from the opposing effects seen in slices lacking astrocyte and neuronal Panx1: the slower frequency seen in slices lacking astrocyte Panx1 could be counteracted by the faster frequency of discharges resultant from the deletion of neuronal Panx1.

Power spectra analysis revealed that Panx1 contribute to increase the power of epileptiform activity induced by KA. The overall power spectra recorded from Panx1-null slices (0.0091 ± 0.0008 mV^2^) exposed to KA was significantly lower than that recorded from Panx1^f/f^ slices (0.0303 ± 0.0035 mV^2^; ANOVA followed by Dunnett’s test, p < 0.0001; Fig. 2J); the decreased overall power of Panx1-null slices was mainly due to a decrease in the delta band (0.5 – 5.0 Hz; Fig. 2H). Compared to Panx1^f/f^ slices, deletion of neuronal or astrocyte Panx1 reduced the overall power spectra of epileptiform discharges (ANOVA followed by Dunnett’s test, p < 0.0001; Fig. 2I, J). These data therefore indicate that deletion of astrocyte or neuronal Panx1 are sufficient to reduce the power of KA-induced hippocampal epileptiform activity to levels seen on Panx1-null slices.

### 3) Both neuronal and astrocyte Panx1 contribute to EEG seizures

To evaluate the impact of Panx1 to acute seizures, we performed EEG recordings in mice injected i.p. with KA (20 mg/kg). Examples of EEG recordings from the four mouse genotypes are shown in Figure 3A. A few min after KA injection, baseline electrical activity changed with the appearance of a high voltage shift (spike, arrows in Fig. 3A) that was followed by short ictal activity duration (seizure; * in Fig. 3A) and then by continuous seizure activity (SE, ! in Fig. 3A). For each mouse, the three recording electrodes (frontal, occipital, and bipolar electrodes; inset in Fig. 3A) displayed the same pattern and duration of ictal events (Supplemetal Figure S5).

**FIGURE 3.**
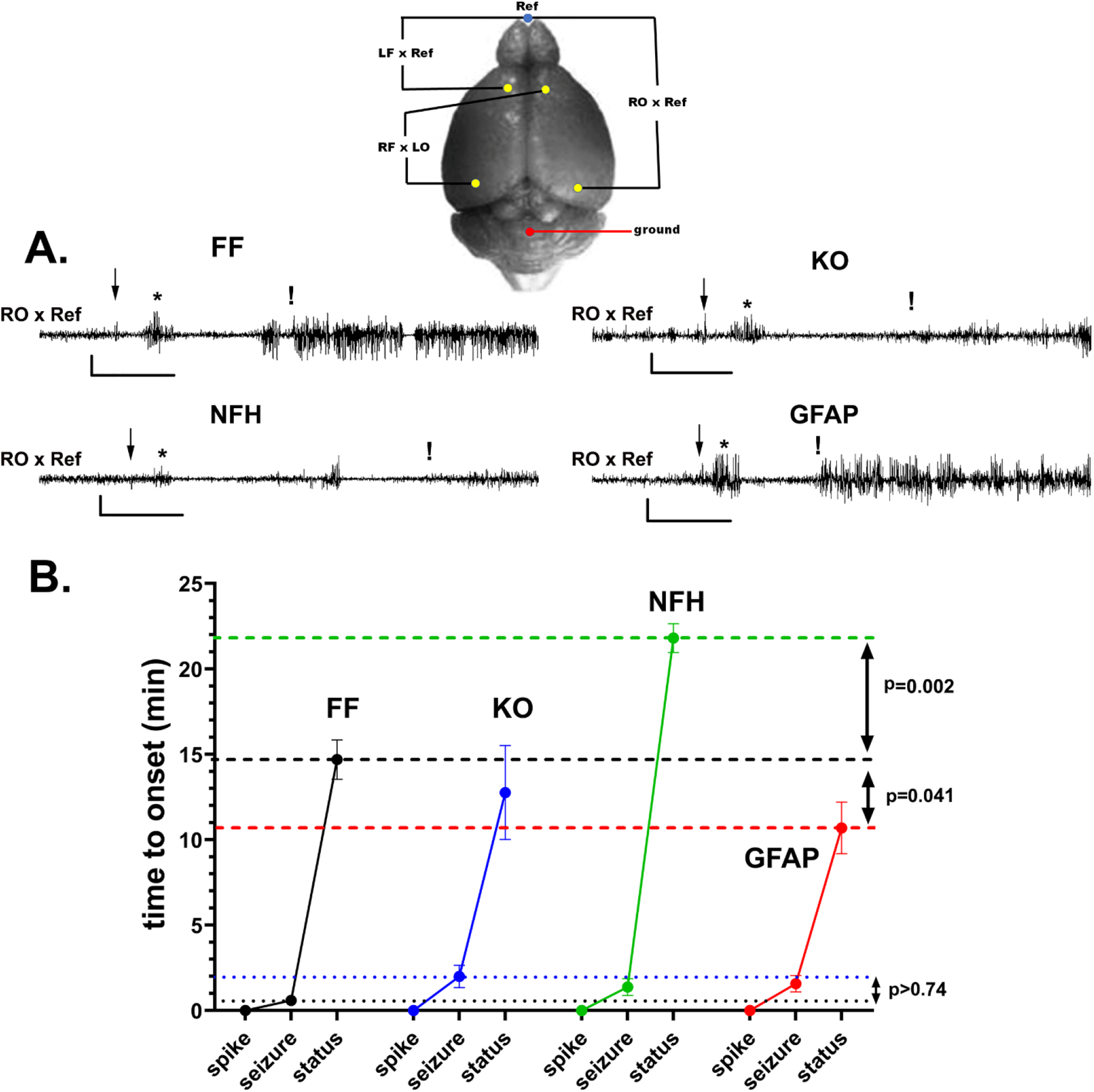
Impact of neuronal and astrocyte Panx1 to acute KA-seizures. **(A)** Representative EEG recordings showing the first spike (arrow), first seizure (*), and status epilepticus (SE) (!) obtained from the occipital cortices of Panx1^f/f^ (FF; n = 7), Panx1-null (KO; n = 5), NFH-Cre:Panx1^f/f^ (NFH; n = 6), and GFAP-Cre:Panx1^f/f^ (GFAP; n = 6) mice. Calibration bars: 300 μV (vertical), 6 min (horizontal). *Inset*: Schematics of electrodes positions on the mouse brain showing the reference electrode (Ref; blue circle) and four recording electrodes (yellow circles), left frontal (LF), right frontal (RF), left occipital (LO), and right occipital (RO); red circle indicates ground position. (**B**) Mean ± s.e.m. values of time to onset of the first seizure and SE following the first spike recorded after intraperitoneal injection of KA (20 mg/kg) in Panx1^f/f^ (black symbols), Panx1-null (blue symbols), NFH-Cre:Panx1^f/f^ (green symbols), and GFAP-Cre:Panx1^f/f^ mice (red symbols). Note the longer time to onset of SE in NFH-Cre:Panx1^f/f^ mice and shorter time to SE onset in GFAP-Cre:Panx1^f/f^ mice. Dotted lines: mean values of the first seizure onset recorded from Panx1^f/f^ (FF) and Panx1-null (KO) mice; dashed lines: mean values of SE onset recorded from Panx1^f/f^ (FF), NFH-Cre:Panx1^f/f^ (NFH), and GFAP-Cre:Panx1^f/f^ (GFAP) mice. The displayed p values are from Dunnett’s multiple comparison tests that followed two-way ANOVA with repeated measures.

To test whether and the extent to which global or cell type specific deletion of Panx1 affected the temporal progression of changes in cortical electrical activity induced by KA, we measured the time to onset of the first seizure (time interval between the first spike and the first seizure) and of SE (interval between the first spike and SE). Two-way ANOVA with repeated measures revealed a significant difference between genotype x ictal events (F_interaction(3, 20)_ = 7.44, p = 0.0014). Post-hoc comparisons indicated no significant difference in seizure onset between Panx1^f/f^ and the other transgenic mice (Dunnett’s multiple comparison, p > 0.74; Fig. 3B). Post-hoc analyses of SE onset indicated no significant differences between Panx1^f/f^ mice (14.69 ± 1.15 min) and Panx1-null mice (12.76 ± 2.75 min; Dunnett’s test, p = 0.53; Fig. 3B); however, compared to Panx1^f/f^ mice, deletion of neuronal Panx1 resulted in a delayed onset of SE (21.80 ± 0.84 min; Dunnett’s test, p=0.002), while deletion of astrocyte Panx1 accelerated SE onset (10.69 ± 1.51 min; Dunnett’s test, p = 0.041; Fig. 3B). Thus, these results suggest that the lack of effect on the SE onset seen in Panx1-null mice may be due to the opposing effects exerted by astrocyte and neuronal Panx1.

Power spectral analysis of EEG ictal activity recorded from the frontal cortex (LF *vs* Ref electrodes, see Fig. 3) indicated no significant difference in overall power spectra between Panx1^f/f^ (2190 ± 83 μV^2^) and Panx1-null mice (2043 ± 150 μV^2^; ANOVA followed by Dunnett’s test, p = 0.59; Fig. 4-A4). Despite the lack of difference in the overall power between Panx1^f/f^ and Panx1-null mice, a significant increase in power in the delta band (0.5 – 5.0 Hz) and decreased power in the beta band (13 – 30 Hz) were detected in the Panx1-null compared to Panx1^f/f^ mice (Fig. 4-A1; ANOVA followed by Dunnett’s test, p<0.0001). In contrast, the overall power spectra obtained from the occipital cortical area (RO *vs* Ref electrode, see Fig. 3) of Panx1-null (1379 ± 109 μV^2^) was significantly lower compared to that of Panx1^f/f^ mice (2028 ± 100 μV^2^; ANOVA followed by Dunnett’s test, p < 0.0001; Fig. 4-B4), particularly in the delta (0.5 – 5.0 Hz) and beta (13 – 30 Hz) bands (Fig. 4-B1).

**FIGURE 4.**
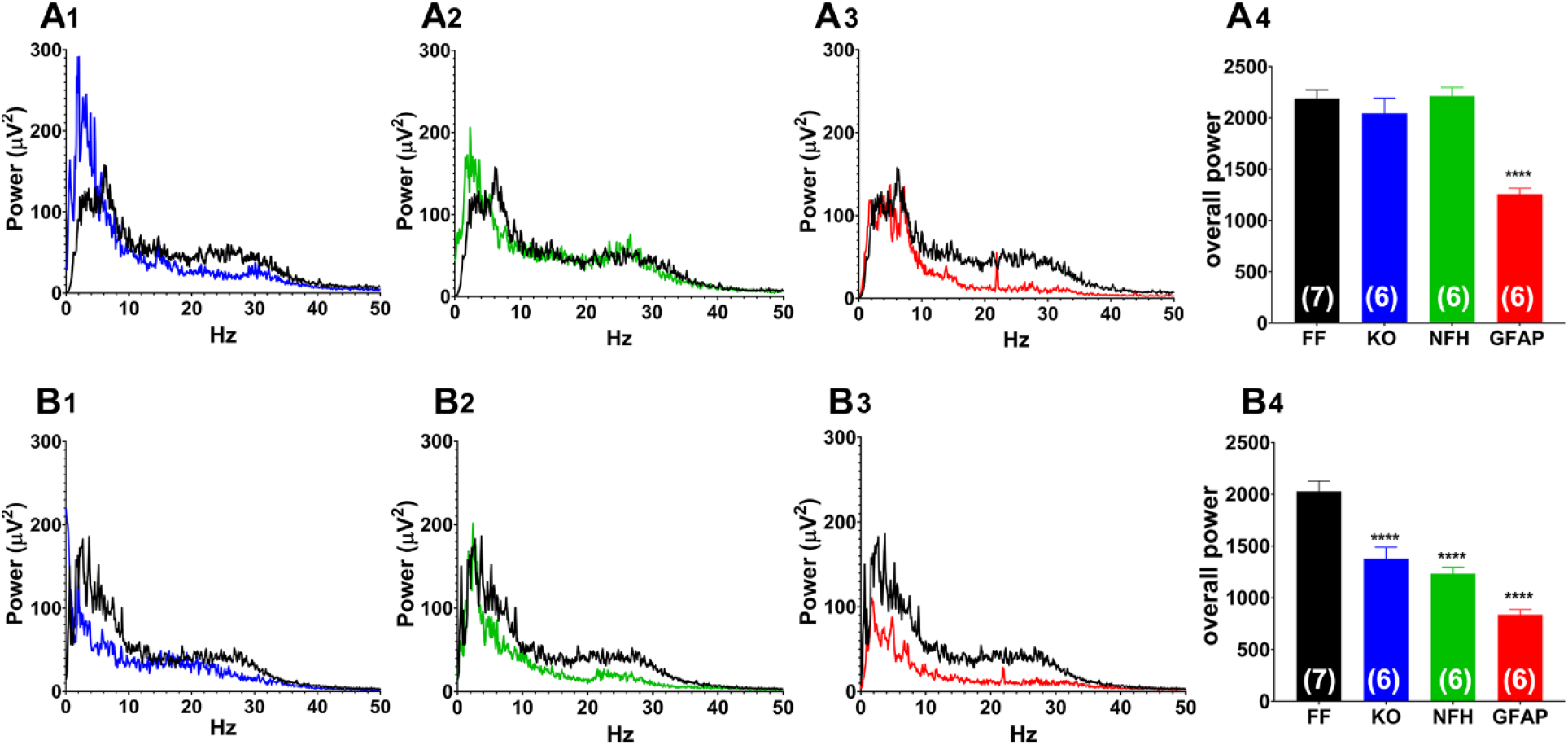
Electroencephalographic (EEG) power spectra obtained from KA-injected mice. (**A-B**) Average power spectra obtained from (**A**) frontal and (**B**) occipital cortices of Panx1^f/f^ (**A1, B1**), Panx1 null (**A2, B2**), NFH-Cre:Panx1^f/f^ (**A3, B3**), and GFAP-Cre:Panx1^f/f^ (**A4, B4**) mice. Mean ± s.e.m. values of the overall power of EEG activity recorded from frontal (**A4**) and occipital (**B4**) cortices obtained from areas under the curves displayed in parts **A1-3** and **B1-3**, respectively. The averaged power spectra and overall power were determined within 0 – 50 Hz frequency range. The number of mice is indicated in the bar histograms. One-way ANOVA followed by Dunnett’s multiple comparison test (****p<0.0001).

To assess the relative contribution of astrocyte and neuronal Panx1 to power spectrum, similar experiments were performed in mice lacking astrocyte or neuronal Panx1. Compared to the overall power recorded from the frontal cortices of Panx1^f/f^ (2190 ± 83 μV^2^), GFAP-Cre:Panx1^f/f^ exhibited lower overall power (1156 ± 57 μV^2^; ANOVA followed by Dunnett’s test, p < 0.0001; Fig. 4-A4), which was mainly due to the reduction in power in all frequencies, except the delta band (0.5 – 5.0 Hz; Fig. 4-A3). Although the overall power recorded from NFH-Cre:Panx1^f/f^ mice did not differ from Panx1^f/f^ (Fig. 4-A4), there was a significant increase in the delta band in mice lacking neuronal Panx1 (Fig. 4-A2; ANOVA followed by Dunnett’s test, p<0.0001). With regards to the occipital cortex, there was a significant decrease in the overall power in mice lacking neuronal and astrocyte Panx1 (NFH-Cre:Panx1^f/f^:1233 ± 64 μV^2^; GFAP-Cre:Panx1^f/f^: 837 ± 50 μV^2^) when compared to Panx1^f/f^ mice (2028 ± 100 μV^2^; ANOVA followed by Dunnett’s test, p<0.0001; Fig. 4.B4); in these two cases, the decreased power was due to attenuations in all frequency bands (Figs 4-B2 and 4-B3).

Thus, power spectral analysis indicates that both astrocyte and neuronal Panx1 contribute to seizures. However, their relative contribution was brain region dependent; while in the occipital cortex, astrocytes and neurons contributed similarly to increase the overall power in all frequency bands, in the frontal cortex, however, deletion of neuronal Panx1 caused an increase in overall power only in the lower frequency band while deletion of astrocyte Panx1 led to a reduction in all frequencies, except the lower band.

## Discussion

Using mice with global and cell type specific deletion of Panx1 in two *in vitro* and a *in vivo* seizure models, we show that both astrocyte and neuronal Panx1 contribute to epileptiform discharges and that their contribution is model and brain region dependent. Under the low-Mg^2+^ condition, global deletion of Panx1 reduced the frequency of discharges, while in the KA-model, global deletion of Panx1 decreased the overall power spectra. In both models, deletion of Panx1 from astrocytes or neurons were not individually sufficient to account for the changes in frequency recorded from Panx1-null slices but were individually sufficient to reduce the power in the KA-model. Finally, our EEG recordings from KA-injected mice revealed a brain region and cell type-dependent effect of Panx1. In the occipital cortex, lack of astrocyte or neuronal Panx1 contributed similarly to reduce the overall power spectra (particularly in delta and beta bands) recorded from Panx1-null. In contrast, in the frontal cortex, Panx1 in these two cell populations had opposing effects (NFH-Cre:Panx1^f/f^: increased delta band power; GFAP-Cre:Panx1^f/f^: reduced beta band power), likely nullifying changes in overall power recorded from Panx1-null mice.

The recorded differences between the epileptiform activity patterns induced by the two ictal inducing agents in Panx1^f/f^ slices are characteristics to the models, with low Mg^2+^ aCSF inducing intermittent bursts of epileptiform activity interrupted by baseline activity and spreading depolarizations (SDs) and KA inducing a short train of epileptiform discharges that is interrupted before KA washout (Engel et al., 2000; Fisher and Alger, 1984; Reid et al., 2008; Velísek et al., 1994). Of note is that epileptiform activity resumed upon washout of KA. This is novel observation that has not been previously reported indicates that the absence of continuous discharges is unlikely due to neuronal cell death, as had been previously suggested (Fisher and Alger, 1984; Routbort et al., 1999; Lopez-Picon et al., 2006).

### Role of Panx1 in seizures

Evidence that Panx1 plays a role in seizure come from studies showing that Panx1 is upregulated in rodent and human epileptic tissues (Jiang et al., 2013; Li et al., 2017; Mylvaganam et al., 2014; Mylvaganam et al., 2010) and that deletion or blockade of Panx1 attenuates KA- and PTZ-convulsant effects (Aquilino et al., 2020; Dossi et al., 2018; Santiago et al., 2011). In contrast, however, Panx1 activation has been shown to attenuate hyperexcitability under hypoglycemic condition (Kawamura et al., 2010; Kawamura et al., 2014) and to decrease seizure susceptibility in the pilocarpine model (Kim and Kang, 2011). These opposing roles of Panx1 on seizures are likely related to diverse downstream signaling pathways triggered by distinct transmitter receptors (see (Scemes and Velíšková, 2017)).

### Panx1 and epileptiform activity

Despite evidence implicating Panx1 in epilepsy, very little is known about the impact that Panx1 has on hippocampal epileptiform activity. A limited number of *in vitro* studies have indicated that Panx1 contributes to increase the frequency of epileptiform discharges and the spectral power induced by two different in vitro models (low-Mg^2+^ and 4-AP) (Thompson et al., 2008; Aquilino et al., 2020). Our data obtained from global Panx1-null hippocampal slices exposed to low-Mg^2+^ condition are in agreement with these previous reports showing a pro-ictal action of Panx1. However, comparing the impact of Panx1 in the two *in vitro* models here utilized, our data also indicate that the mode of action of Panx1 is model dependent. While under the low-Mg^2+^ condition we found reduced frequency of epileptiform discharges without changes in power spectra, the opposite was obtained for hippocampal slices exposed to KA (no changes in frequency but decreased power). Such model-dependent effect is also evidenced in preparations in which Panx1 was deleted in a cell-type specific manner. Differently from the low-Mg^2+^ condition, where neither deletion of Panx1 from astrocytes or from neurons resulted in the decreased frequency of discharges recorded in Panx1-null slices, in the KA-model Panx1 deletion from neurons or astrocyte had opposing effects on the frequency of discharges but were similarly sufficient to reduce the power spectra. It is possible that such model-dependent effects of Panx1 are related to different mechanisms by which Panx1 can be activated (Lohman et al., 2019; Scemes and Spray, 2012; Silverman et al., 2009; Weilinger et al., 2016), which could differently impact Panx1-mediated purinergic signaling. It has been reported that depending on the stimuli, distinct conductive states of Panx1 can be induced, a high conductive non-selective state permeable to ATP and cationic fluorescent dyes and a low conductive state that is anion selective but not permeable to ATP (Wang et al., 2014). With regards to glutamate-mediated Panx1 activation, both ionotropic and metabotropic receptors can lead to opening of the high conductive state of Panx1 as seen by the increased release of ATP and influx of fluorescent dyes in hippocampal slices (Lopatář et al., 2015; Santiago et al., 2011; Thompson et al., 2008). This occurs despite the different downstream signaling pathways induced by glutamate receptor subtypes. While opening of Panx1 channels can be induced by NMDA receptor-mediated src kinase phosphorylation at Y308 (Lohman et al., 2019; Weilinger et al., 2016), opening of Panx1 channels due to intracellular Ca^2+^ rise (Locovei et al., 2006) can be induced by mGluR5 (Lopatář et al., 2015) and likely by KA.

In addition to the mode of Panx1 activation, it is also possible that the differential expression of purinergic (P2 and P1) receptors among the cell types in different brain regions (Burnstock et al., 2011; Köles et al., 2011) may contribute to distinct network outputs when Panx1 is deleted. Regarding P2 receptors, it has been shown that Panx1-mediated ATP release acting on P2Y1 receptors contribute to mGluR5-mediated bursting activity of CA3 but not of CA1 hippocampal subfield (Lopatář et al., 2015). Under low-Mg^2+^ condition, Panx1-mediated ATP release increase ictal activity in human cortical tissues (Dossi et al., 2018) but minimally affects epileptiform discharges in rodent hippocampal slices (Lopatář et al., 2011). In contrast to the effects of P2 receptors, the action of adenosine on P1 receptors is well known to have anti-convulsant effects (Boison, 2012; Boison, 2016; Cieślak et al., 2017; Dale and Frenguelli, 2009; Masino et al., 2014). This inhibitory effect of adenosine relates to its action on the high affinity A1 receptors which are abundantly expressed in the hippocampus and neocortex, brain areas prone to seizures (Masino et al., 2014). Although the pro-convulsant effects have been reported for other adenosine receptor subtypes (A2 and A3), they are less prominent than the effects of adenosine on A1 receptors given their very low expression levels and restricted brain distribution (Masino et al., 2014).

Thus, the model-dependent effect of Panx1 on epileptiform discharges found in our study can be explained by the combined effects of activation modes of Panx1, the distribution of P2 and P1 receptors, and the output generated by neuron-glia interactions. Further studies aimed to investigate the contribution of astrocyte and neuronal Panx1 to purinergic mediated signaling in hyperexcitability are necessary to provide more detailed mechanistic insight.

### Panx1 and acute seizures

The few studies that have addressed the question of whether Panx1 plays a role during the initial phases of acute seizures indicated that, except for the pilocarpine model, deletion or blockade of Panx1 have protective effects (Aquilino et al., 2020; Kim and Kang, 2011; Santiago et al., 2011). Here we show using EEG recordings from Panx1-null mice injected with KA that Panx1 contributed to increase the power spectra, particularly in the beta band, but has no effect on the onsets of the first seizure or status epilepticus. A recent study characterizing behavioral seizures and the EEG correlates induced by KA, indicated that when the seizures progressed from non-convulsive (Racine stages 1 and 2) to convulsive (Racine stages 3–5), the power in different bands varied, with higher delta and theta powers appearing during stage-2, and increased beta and gamma powers during the progression from non-convulsive to convulsive seizures (Sharma et al., 2018). Thus, the reduced beta band power recorded from the frontal and occipital cortices of Panx1-null suggests that the progression of non-convulsive to convulsive seizures is “attenuated” in these mice. Such interpretation agrees with our previous studies showing that deletion or blockade of Panx1 improves seizure scores (Santiago et al., 2011). Similarly, Panx1 blockade or deletion reduced the duration of afterdischarges and prevented electrical kindling as well as reduced seizure severity and death following systemic PTZ administration (Aquilino et al., 2020).

Interestingly, we found using power spectral analysis, brain region and cell-type dependent effects of Panx1 that have not been previously reported. In the occipital cortex, deletion of neuronal or astrocyte Panx1 contribute to decrease the overall power, mainly reducing delta and beta bands; in contrast, in the frontal cortex deletion of neuronal Panx1 led to increased theta band while deletion of astrocyte Panx1 led to decreased delta and beta bands. Although the reason for these differences is unknown, it is possible that it may result from a differential expression levels of Panx1 and/or distinct neuron-to astrocyte ratios in these two cortical areas. It has been reported that Panx1 expression levels vary in distinct CNS areas with higher expression in hippocampus and lower in cerebellum (Hanstein et al., 2013). Glia-to-neuron ratio has been found to be higher in the cortex than in cerebellum (Herculano-Houzel, 2014).

Regarding the onset to SE recorded here in the Panx1-null mice, it is possible that the lack of impact on SE onset in the null genotype results from the opposing effects recorded from mice lacking neuronal Panx1 (delayed SE onset) and from those lacking Panx1 in astrocytes (accelerated onset). Nevertheless, these results are in agreement with our previous study showing delayed onset of bilateral front limb clonus in NFH-Cre:Panx1^f/f^ and delayed onset in GFAP-Cre:Panx1^f/f^ (Scemes et al., 2019). The present and previous studies indicate that neuronal Panx1 substantially promotes excitability while astrocyte Panx1 exerts inhibitory action instead. The mechanism underlying these opposing effects on the progression of KA-induced seizures is related to the increased expression levels of adenosine kinase (ADK) in astrocytes as measured in GFAP-Cre:Panx1^f/f^ brains (Scemes et al., 2019). This increased level of ADK, an enzyme that by phosphorylating intracellular adenosine to AMP (Rotllan and Miras Portugal, 1985), can together with the action of the equilibrative nucleoside transporters (ENTs) favor the flux of adenosine from the extracellular to the astrocyte compartment (Parkinson et al., 2011; Lovatt et al., 2012; Wall and Dale, 2013; Boswell-Casteel and Hays, 2017; Griffith and Jarvis, 1996; Zhang et al., 2011). Consequently, the inhibitory action of adenosine mediated by A1 receptors would be reduced, favoring a faster progression of seizures in these mice.

In conclusion, we show here using *in vitro* and *in vivo* seizure models that both astrocyte and neuronal Panx1 contribute to ictal activity and that their contribution is model and brain region dependent.

## Acknowledgemets

This work was supported by the National Institute of Neurological Disorders and Stroke, National Institutes of Health (5R01-NS-092786 to ES and JV). The authors acknowledge the contribution of Mr. Jian Pan for his technical support and for maintaining mouse colonies.

