## Supplementary material for "The contribution of astrocyte and neuronal Panx1 to seizures is model and brain region dependent": SUPPLENATL FIGURES S1-S5

**SUPPLEMENTAL MATERIAL**

**FIGURE S1**. *Restoration of baseline activity following low-Mg^2+^ washout*. Example of extracellular field potential recording obtained from the CA1 region of a Panx1-null hippocampal slice showing the development of epileptiform activity following exposure to Mg^2+^-free aCSF (arrow) and reestablishment of baseline after washout (arrowhead) with normal aCSF. Calibration: 10mV (vertical), 20 min (horizontal). Note the eventual and marked decrease of discharges following washout.

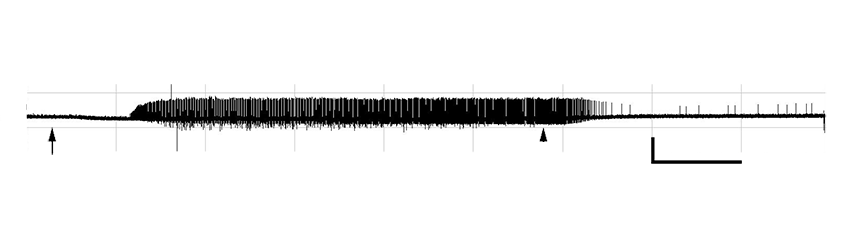

**FIGURE S2**. *Spreading depolarization induced by low-Mg^2+^ aCSF*. Examples extracellular field potential recordings obtained from CA1 regions of Panx1^f/f^ (FF, top) and Panx1-null (KO, bottom) hippocampal slices showing large (~-10 to -20 mV) long lasting (up to 1 min) negative deflections from baseline, characteristic of spreading depolarizations (SDs). Arrows indicates low-Mg^2+^ application; *SDs. Calibration bars: 20 mV (vertical), 20 min (horizontal).

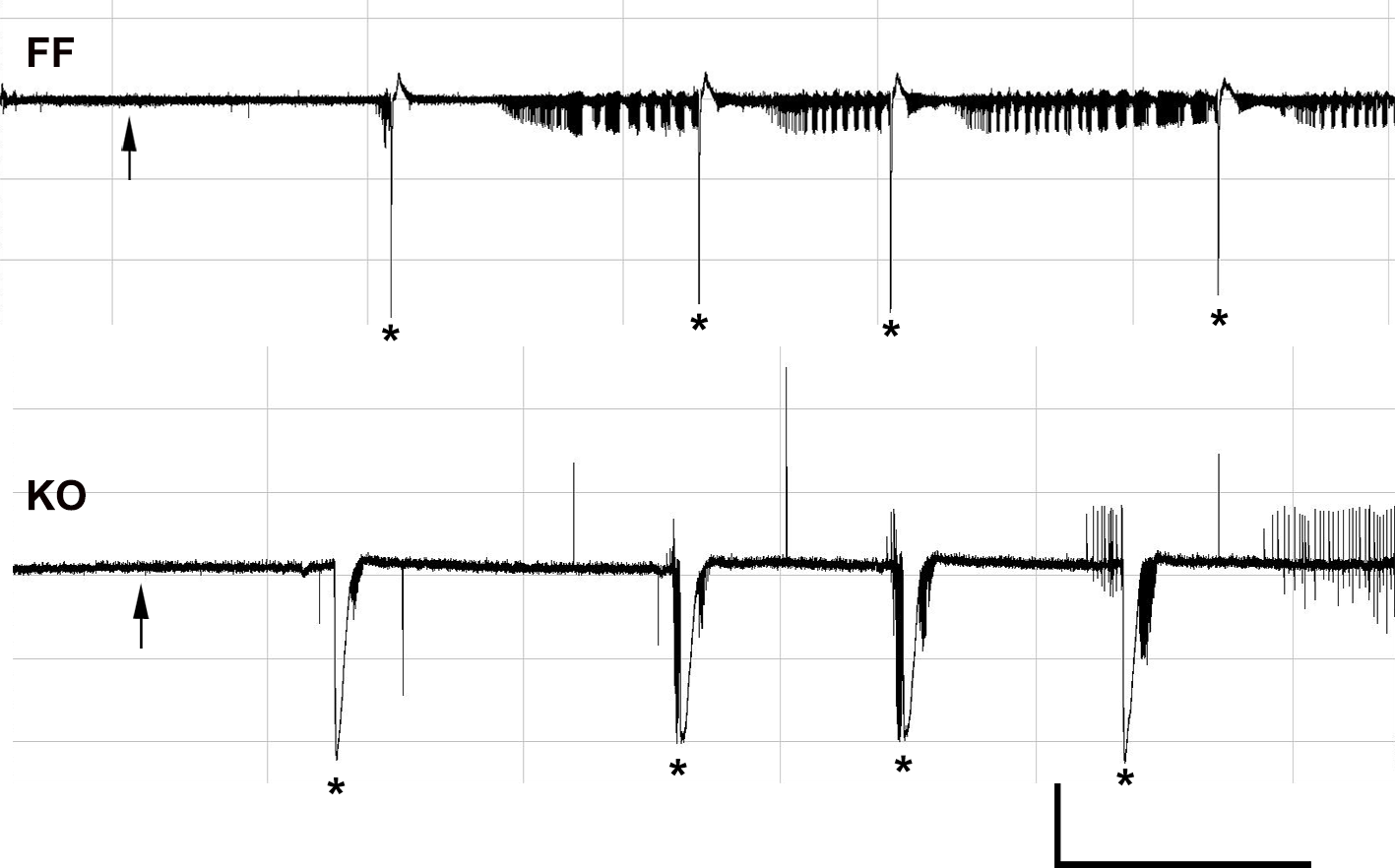

**FIGURE S3.** *Patterns of epileptiform discharges***.** Representative extracellular field potential recordings obtained from CA1 regions of NFH-Cre:Panx1^f/f^ (NFH, top) and GFAP-Cre:Panx1^f/f^ (GFAP, bottom) mice hippocampal slices following application of low-Mg^2+^ aCSF (indicated by arrows) showing the burst-like pattern of epileptiform discharges. Calibration: 10 mV (vertical), 20 min (horizontal).

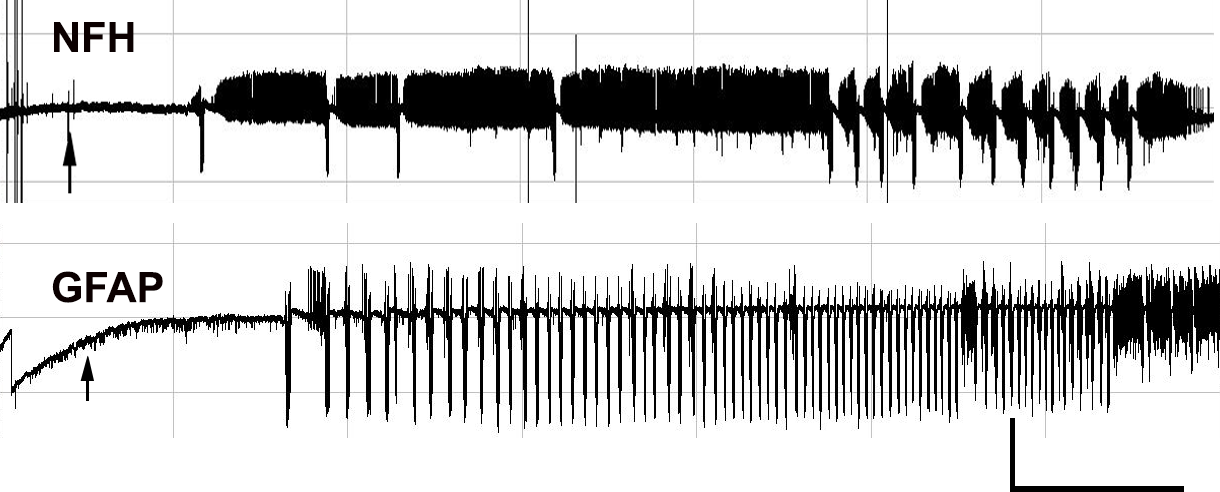

**FIGURE S4.** *Washout of KA does not restore baseline activity***.** Example extracellular field potential recording obtained from the CA1 region of a Panx1-null mouse hippocampal slice showing the epileptiform activity induced by kainic acid (KA) exposure (arrow). After KA washout (arrowhead), epileptiform activity resumes. Calibration: 5.0 mV (vertical), 20 min (horizontal).

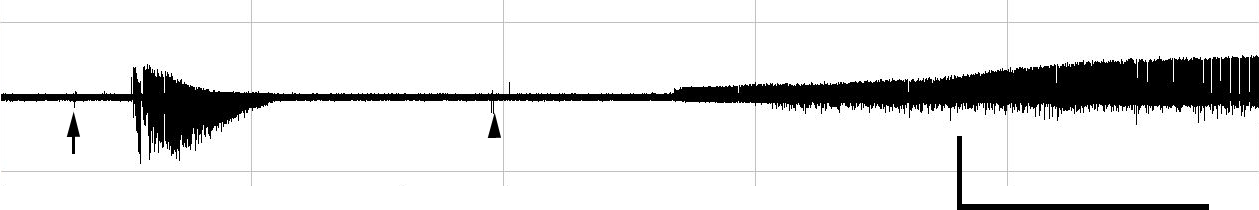

**FIGURE S5.** Example of EEG activity detected by three electrode pairs *LF x Ref*, *RO x Ref*, and *RF x LO* placed on the cortex of a Panx1^f/f^ mouse injected with KA (20mg/kg) showing the time to the first spike (arrows), first seizure (*) and SE onset (!). LF: left frontal electrode; RO: right occipital electrode; RF right frontal electrode; LO: left occipital electrode; Ref: reference electrode. Calibration: 400 µV (vertical), 5 min (horizontal).

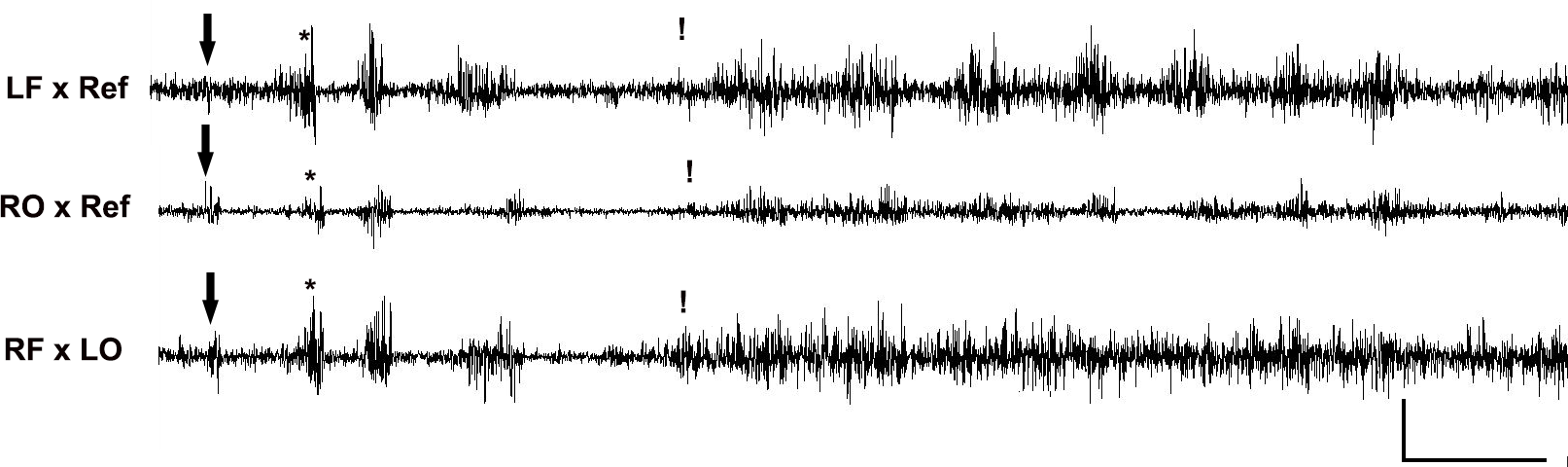
